# Clover-like object vector representations support human spatial cognition

**DOI:** 10.1101/2025.04.18.649400

**Authors:** Marcia Bécu, Ivan Markel Krasovec, Pearl Saldanha, Jonathan R. Whitlock, Christian F. Doeller

## Abstract

Vector-based spatial coding has been demonstrated in rodents, yet its neural basis in humans and its contribution to navigation remain poorly understood. Using a novel spatial updating task, we show that egocentric directional signals peak when objects are located behind the navigator, while distance signals emerge only when objects are out of view, suggesting a mnemonic role for vision-independent spatial mapping. Allocentric signals form a clover-shaped, four-axis pattern aligned to visual features, with improved navigation accuracy along its axes. Parallel rodent recordings further show the clover pattern arises from averaged activity of allocentric vectorial neurons, suggesting conserved cross-species mechanisms. Together, our findings uncover vector-based representations in the human brain, potentially serving as a neural reference axis to anchor objects to internal maps and support navigation.

---

Vector-responsive cells represent the presence of salient cues, objects or boundaries, at specific directions and distances relative to an agent (*1–7*) and have been described in the rodent brain in an egocentric (self-centered) frame of reference (eg. in parietal, retrosplenial, postrhinal cortex and striatum) (*3, 8, 9*) and in an allocentric frame of reference (eg. in entorhinal cortex, subiculum and hippocampus) (*2, 4, 5*). Combined with self location representation (place cells) (*10*), path integration (grid cells) (*11*) and heading representation (head-direction cells) (*12*), object- and boundary-vector codes theoretically allow the mapping of geometric relationships between the self and the external world through self-motion, even when cues are out of sight. Yet, a direct assessment of vector representation functions and their precise role in supporting navigation remains lacking, especially in humans.

If all distances and directions were equally represented in the brain, it would theoretically be impossible to detect any macroscopic vectorial signal in the human brain using functional magnetic resonance imaging (fMRI) (*13*). However, evidence shows that vector-sensitive receptive fields cover space non-uniformly: for instance, populations of egocentric boundary cells in the retrosplenial cortex show a preferential tuning to short distance ranges and directions behind the animal (*3*), while populations of allocentric object-vector cells in medial entorhinal cortex show sparser representations of distances farther from an object (*2*).

Based on this assumption, the present study investigated how allocentric and egocentric direction and distance to objects are represented in the human brain. We used a novel spatial updating task, in which participants pointed to an intramaze object following periods of self-motion, providing a direct behavioral measure of object location representation. Based on the above rodent studies and studies in humans (*14, 15*), the locations expected to represent vectorial information include the posterior parietal cortex (PPC), retrosplenial cortex (RSC), parahippocampal cortex (PHC), entorhinal cortex (EC) and hippocampus (HP). We also sought to characterize the neural population tuning response in relation to object distance/direction, and verify the functional relevance of these codes for human spatial orientation and navigation.

## Mapping directional and distance tuning

We used 7 Tesla fMRI to monitor the brain activity of 47 young participants navigating a virtual arena surrounded by distal landmarks (a mountain scenery and a castle) and with a large object (a column) situated in the northeast quadrant (Fig. 1A-B). During the task, participants alternate between actively navigating to start locations scattered around the object and being passively transported along predefined trajectories. Throughout the passive transportation the object is invisible, but participants must keep track of its location and point back to it (Fig. 1C-D). The current object vector experienced from a given start location is updated by self-motion (experienced while being passively moved along a given trajectory) in order to be able to accurately point back at the object at the end of the trajectory. Start and end locations of the trajectories were rigorously controlled in order to ensure full coverage of all directions and distances to the intramaze object (see Methods and fig. S1A-C). Responses occurred either from an egocentric perspective (i.e. egocentric condition: ”the object is ahead/behind/left/right of me at the end of the trajectory”) or from an allocentric perspective (i.e. allocentric condition: ”I am north/east/south/west of the object at the end of the trajectory”). While both egocentric and allocentric representations theoretically coexist during spatial navigation, the two conditions and trajectories were designed to provide a behavioral readout of spatial updating performance at different directions and distances to the object and along the two frames of reference (allocentric and egocentric). Two categories of fMRI models were estimated: 1) parametric modulation analyses were used as ’localization’ models to identify voxels in the brain which activity varied with *(i)* the egocentric direction to the object, *(ii)* the allocentric direction to the object and *(iii)* the distance to the object, whereas 2) tuning models discretized the data into evenly-spaced bins in order to characterize the BOLD response variation to changing self-object relationships (see Methods for model specifics). Group-level peak activity in the parametric modulation models served as *a priori* locations for the tuning model’s estimation.

**Figure 1:**
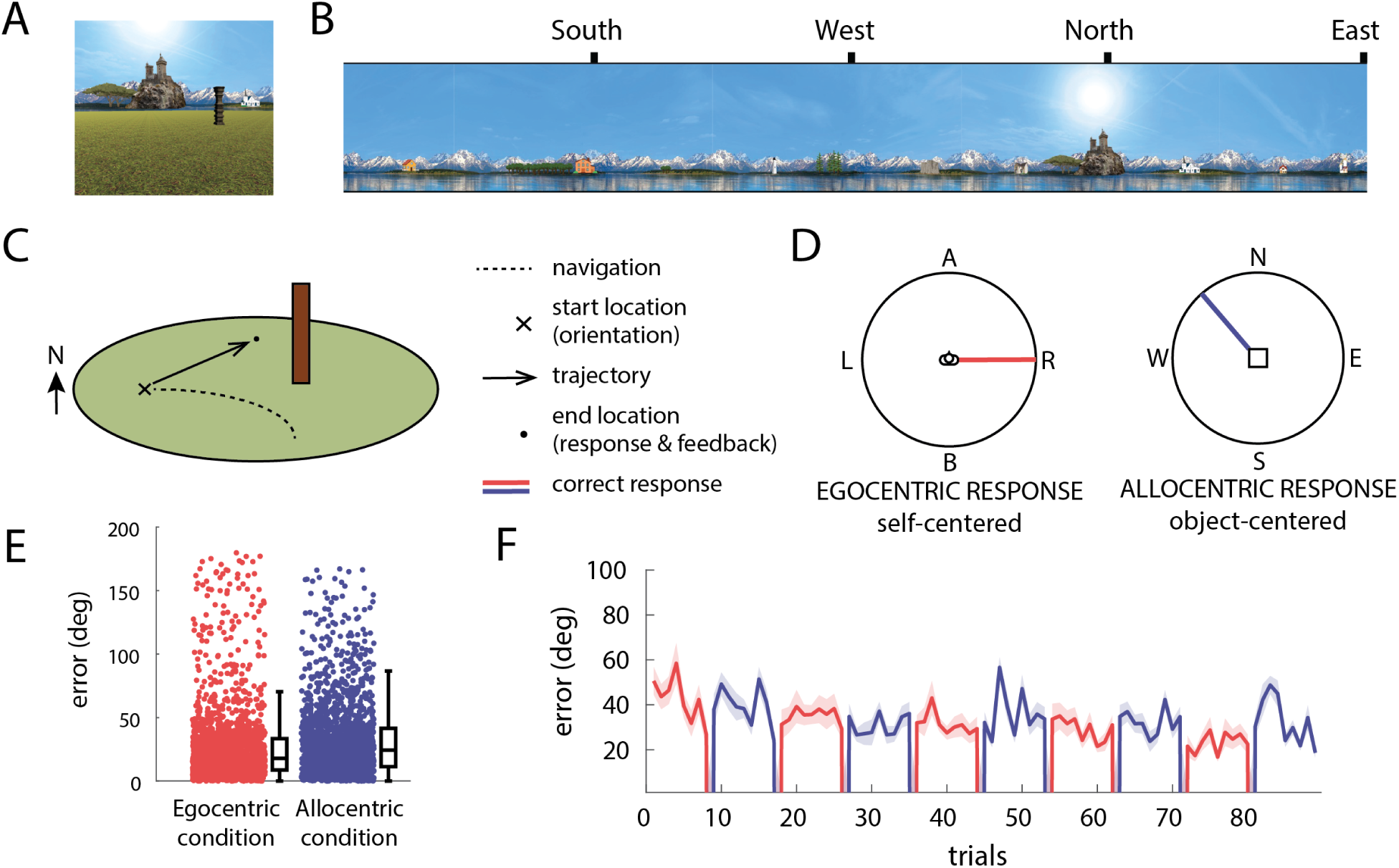
Spatial navigation task in virtual reality (VR) (A) First-person view of the environment with the intramaze object and (B) array of distal landmarks projected at infinity in the background. The main distal landmark (i.e. castle) situated in the North. (C-D) Trial sequence: Participants actively navigate to a start location and freely explore. After a key press, the object disappears and they are passively moved along a fixed path while tracking the object’s location mentally. At the end point, they indicate the object’s position using an egocentric or allocentric response. Responses (in degrees) are followed by performance-based feedback. (E) Pointing error in degrees accumulated across all blocks and participants and (F) along the 10 blocks of egocentric (red) and allocentric (purple) trials

## Spatial updating performance and gaze pattern

Pointing error in the allocentric condition was slightly higher than the egocentric condition (𝐹(_46,1_) = 7.07, 𝑝 = 0.011, generalized eta-squared (ges): 𝜂_g_^2^ = 0.027, Fig. 1E-F). Two-way ANOVAs show a significant effect of the direction at which the trajectory lies in both the egocentric condition (direction: 𝐹_(1.71,68.45)_ = 16.002, *p* < 0.0001, 𝜂_g_^2^ = 0.066, distance: 𝐹_(2,80)_ = 0.09, *p* = 0.9, 𝜂_g_^2^ = 0.0002, interaction: 𝐹_(4,160)_ = 0.36, *p* = 0.8, 𝜂_g_^2^ = 0.002) and allocentric condition (direction: 𝐹_(2,78)_ = 24.19, *p* < 0.0001, 𝜂_g_^2^ = 0.094, distance: 𝐹_(2,78)_ = 2.14, *p* = 0.12, 𝜂_g_^2^ = 0.006, interaction: 𝐹_(4,156)_ = 1.54, *p* = 0.19, 𝜂_g_^2^ = 0.008, fig. S1D), with trajectories going towards the object associated with lower pointing performance. This analysis was performed for n=41/40 since there were missing observations in 6/7 participants in the egocentric or allocentric condition, respectively.

To understand which cues participants used during the task, we estimated gaze dwell times during the orientation phase, where participants had unlimited time to look around and orient themselves at the start location. We binned gaze locations to three areas of interest in the environment: the background (where distal landmarks are), the object and the ground (fig. S1E). We found that participants spent significantly more time looking at the distal landmarks (Wilcoxon signed-rank 𝑊 = 54, *p* < 0.0001, 𝑟 = −0.74), particularly at the northern-most landmark (the castle, see. fig. S1F) in the allocentric condition compared to the egocentric condition, while the time spent looking at the object (the column) was similar in both conditions (𝑊 = 415, *p* = 0.34, 𝑟 = 0.16). In contrast, participants spent more time looking at the ground, close to the object location (Fig. S1F) during the egocentric condition (𝑊 = 568, *p* = 0.0002, 𝑟 = 0.60).

## Body-axis directional tuning of egocentric representations

We first estimated the localization model with a parametric modulation analysis where direction to the column was mapped to the interval [-1,1], where -1 corresponds to the object being behind the participant and 1 to the object being in front of the participant. We identified three key effects in our ROIs: a positive modulation in the PPC (coordinates: [48,-50,44], Brodmann area 39), and negative modulations in both the RSC (coordinates: [20,-51,8], Brodmann area 23) and the PPC (coordinates: [21,-42,-8], Brodmann area 36) (Fig. 2A). To better understand how the activity varied with changing self-object relationships, we further discretized the observed egocentric directions into 8 evenly-spaced bins (tuning model specified in 2B-C). Here, we found higher signal in the PPC when the object was straight ahead of the participant, while in the RSC and the PHC, activity peaked when the object was behind the participant (bins 1 and 8) (Fig. 2D-G).

**Figure 2:**
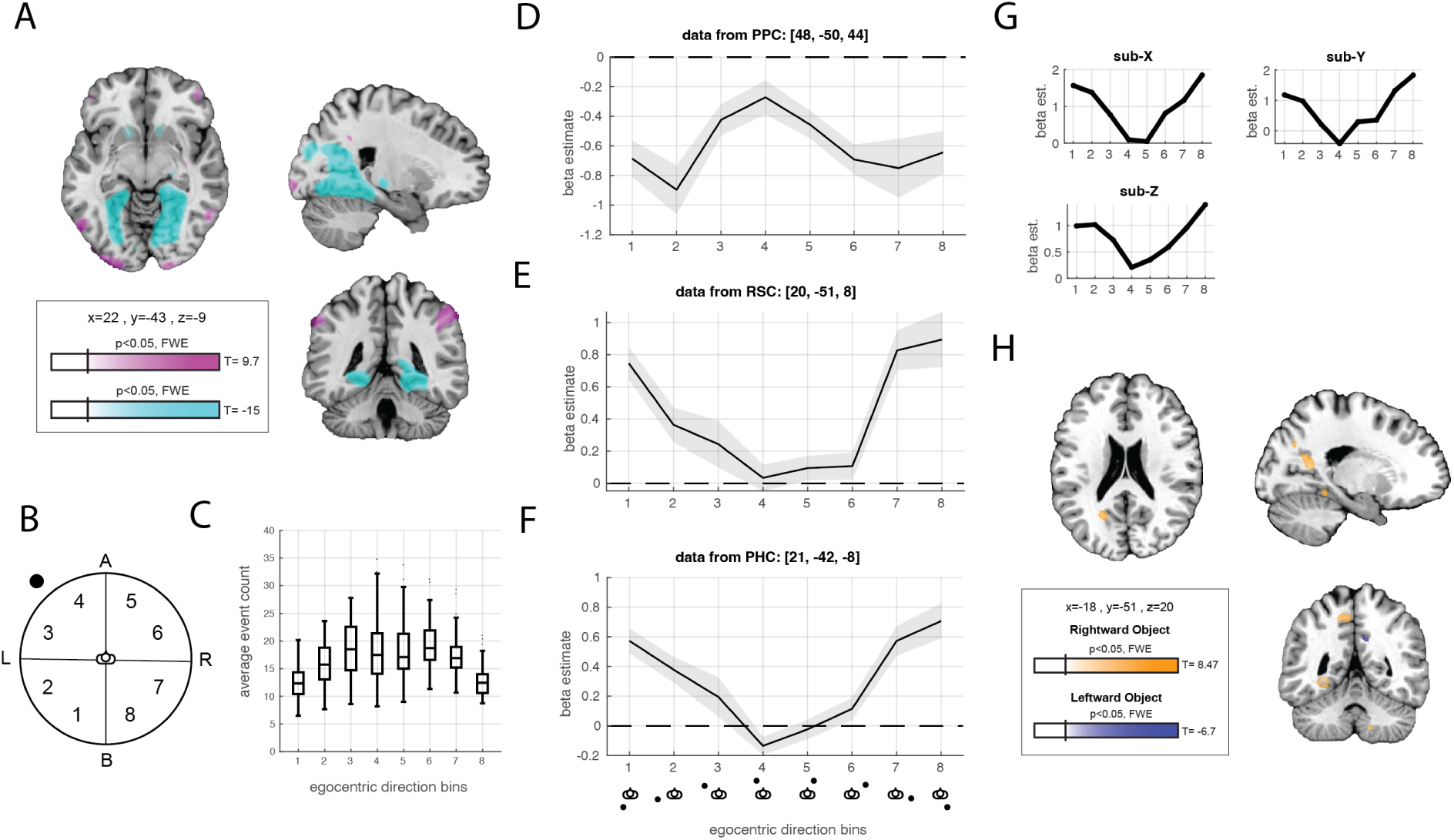
Egocentric directional modulation of the fMRI signal. (A) Localization model: whole brain effect showing a positive/negative parametric modulation of activity by egocentric direction in pink/blue, respectively. Three peaks identified in our regions of interest: posterior parietal cortex (PPC), retrosplenial cortex (RSC), parahippocampal cortex (PHC). Maximum, minimum T-scores, and MNI slice location. FWE=family-wise error correction. (B-C) Egocentric directions (ahead, behind, left, right) were discretized into 8 equidistant bins used to estimate the tuning model in a 2-voxel-radius sphere centered on previously identified peaks. (C) Number of events per bin, with the median, 25th and 75th percentiles (black box), 1.5 x interquartile range (whiskers), and outliers (dots). (D-F) Median tuning curves for different egocentric directions bins. Shaded areas indicate the between-subject standard error of the mean (SEM). (G) Individual tuning curves in RHC and PHC shown as best examples. (H) Hemispheric lateralization model: whole brain parametric modulation with positive/rightward object and negative/leftward object modulation in yellow and blue, respectively. On the slice, the left hemisphere appears on the left, following neurological convention.

A one-way ANOVA with egocentric direction bins as a within-subjects factor showed a significant omnibus effect in the three ROIs: PPC (𝐹_(5.64,208.65)_ = 8.65, *p* < 0.001, 𝜂_g_^2^ = 0.093), RSC (𝐹_(4.51,175.91)_ = 35.21, *p* < 0.001, 𝜂_g_^2^ = 0.25) and PHC (𝐹_(4.66,172.57)_ = 44.44, *p* < 0.0001, 𝜂_g_^2^ = 0.30). Post-hoc polynomial contrast showed a significant quadratic and, to a lesser extent, quartic tendencies in the three ROIs: in PPC (linear: 𝑡_(37)_ = −0.27, *p* = 0.79; quadratic: 𝑡_(37)_ = −7, *p* < 0.0001; cubic: 𝑡_(37)_ = 1.09, *p* = 0.28; quartic: 𝑡_(37)_ = 3.13, *p* = 0.003; 9 outliers removed), RSC (linear: 𝑡_(39)_ = 2.25, *p* = 0.03, does not survive Bonferroni correction; quadratic: 𝑡_(39)_ = 11.81, *p* < 0.0001; cubic: 𝑡_(39)_ = −0.37, *p* = 0.71; quartic: 𝑡_(39)_ = −3.53, *p* = 0.001; 7 outliers removed) and PHC (linear: 𝑡_(37)_ = 2.49, *p* = 0.01, does not survive Bonferroni correction; quadratic: 𝑡_(1,37)_ = 11.83, *p* < 0.0001; cubic: 𝑡_(37)_ = 0.91, *p* = 0.36; quartic: 𝑡_(37)_ = −7.04, *p* < 0.0001; 9 outliers removed).

To better understand if egocentric direction modulation gave any advantage to navigation, we correlated the average parametric modulation from the localization model (Fig. 2A) with task performance and two spatial cognitive tests performed at the end of the experiment (the Perspective Taking test and 4 Mountains test, fig. S2A-C). Generally, we found the error (expressed as a z-score) to be lower, thus performance was better in people showing the strongest modulation in PPC, but this correlation did not survive multiple comparison correction (egocentric: 𝑟 = −0.34, *p* = 0.02; allocentric: 𝑟 = −0.26, *p* = 0.074, fig. S2A). No other correlation in RSC and PHC were significant (fig. S2B-C).

Finally, since rodent data indicates a contralateral bias in representing egocentric vector to boundaries (*3*) (that is, left RSC represents contralateral directions and vice-versa), we tested for hemispheric lateralization with right/leftward object being mapped as 1/-1 (see Fig. 2I and caption). We indeed observed two contralateral peaks of activation in RSC, with the left hemisphere coding for when the object was situated to the right of the subject (coordinates: [-18, -58, 12], Brodmann area 23), and the right hemisphere coding for when the object was situated to the left of the subject (coordinates: [11, -50, 32], Brodmann area 23). Additional peaks were found in the left hemisphere (in parietal cortex at coordinates: [-6, -50, 50], Brodmann area 7 and close to PHC at coordinates [-28, -49, -6], Brodmann area 19).

## Clover-like directional tuning of allocentric representations

For allocentric directions, we initially mapped north/south directions as 1/-1, and found 2 clusters of direction-modulated activity in our areas of interest: RSC (coordinates: [20,-56,13], Brodmann area 23) and PHC (coordinates: [21,-35,-12], Brodmann area 36). When estimating the tuning models with 8 allocentric direction bins, we discovered that activity in those areas increased at three or four seemingly orthogonal directions around the object (bins 2/4/6/8, Fig. 3F). This pattern of activity (peaks in bins 2/4/6/8, troughs in bins 1/3/5/7) was observed in most participants (see individual examples on Fig. 3H). Based on this observation, we ran an additional localization model that mapped bins 2/4/6/8 as 1 and 1/3/5/7 as -1 (see Materials and Methods). This model confirmed our clusters in the RSC and PHC (Fig. 3A, positive modulation) and identified a third peak in EC, particularly in the posterior-medial section (pmEC, Fig. 3B, negative modulation). When estimating the tuning model in pmEC (Fig. 3C-D), we observed that, contrary to RSC and PHC, activity in EC was higher in bins 1/3/5/7 (light areas on Fig. 3E, left). Nevertheless, the pattern in EC was similar, with the BOLD signal being higher around three or four directions to the object. One-way ANOVAs with directional bins as a within-subjects factor confirmed a significant omnibus effect in all three ROIs (RSC: 𝐹_(3.98,163.28)_ = 6.37, *p* < 0.001, 𝜂_g_^2^ = 0.036; PHC: 𝐹_(4.53,167.75)_ = 4.46, *p* < 0.001, 𝜂_g_^2^ = 0.032; EC: 𝐹_(5.22,177.36)_ = 5.13, *p* = 0.025, 𝜂_g_^2^ = 0.023) and a post-hoc contrast confirmed that the activity in bins 2/4/6/8 significantly differed from activity in bins 1/3/5/7 (RSC: 𝑡_(41)_ = −4.442, *p* < 0.0001; 5 outliers removed; PHC: 𝑡_(37)_ = −4.446, *p* < .0001; 9 outliers removed; EC: 𝑡_(34)_ = 2.43, *p* = 0.02; 12 outliers removed).

**Figure 3:**
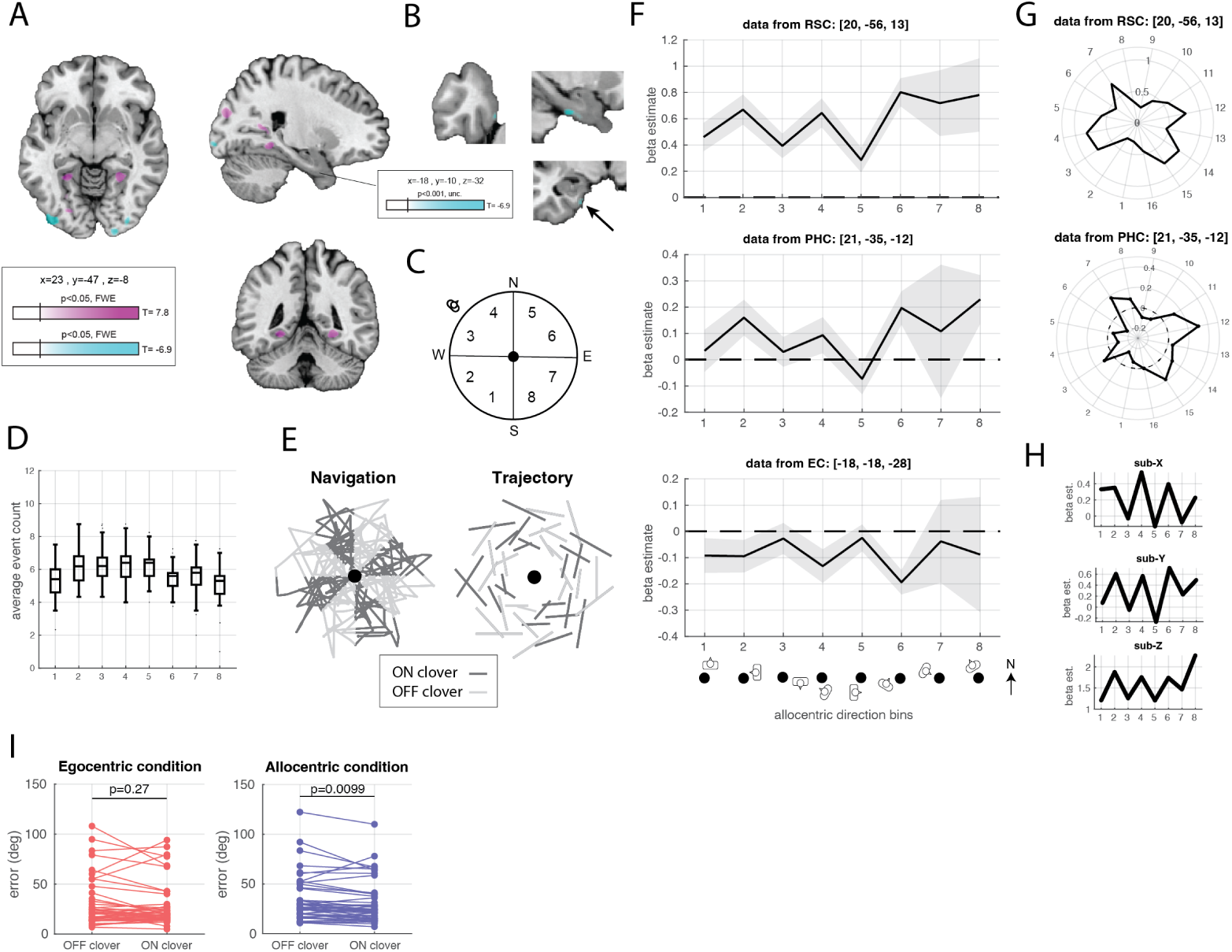
Allocentric directional modulation of the fMRI signal. (A) Localization model: whole brain effect with positive/negative parametric modulation of activity by allocentric direction in pink/blue, respectively. Three peaks identified in our regions of interest: retrosplenial cortex (RSC), parahippocampal cortex (PHC) and entorhinal cortex (EC, though at uncorrected threshold). Maximum, minimum T-scores, and MNI slice location shown. FWE=family-wise error correction. unc=uncorrected threshold *p* < 0.001. Data estimated with the pure clover model. (C-D) Allocentric directions discretized into 8 equidistant bins to estimate the tuning model. (D) Number of events for each bin with median, 25th and 75th percentiles (black box), 1.5 x interquartile range (whiskers), and outliers (dots). (E) Individual example of navigation (left) and trajectory (right) paths lying on (dark gray) or off (light gray) clover axes (F) Median tuning curves across 8 allocentric directions bins. Shaded areas show SEM. (G) Median tuning curves across 16 allocentric direction bins. Outlier data points were removed before calculating the group median. (H) Individual tuning curves for the 8 shown as best examples. (I) Pointing error (in degrees) on and off the clover axes in the egocentric (left) and allocentric (right) conditions. Errors averaged across bins.

To better understand this pattern of neural responses, we ran a more detailed tuning model with 16 bins representing 16 discretized allocentric directions relative to the object (Fig. 3G). The 3/4 peaks in RSC and PHC, although not entirely orthogonal to each other, formed a clover-like pattern of activity around the object. In reference to the main landmark situated in the North (between bins 8 and 9 on Fig. 3G), the closest peak in directional bin 7 was located 33.75◦ away from the main North axis defined by the environment.

When correlating the strength of the parametric modulation with task performance, we found positive correlations whereby stronger clover-like modulation was associated with being better at pointing back at the object, although statistics did not survive multiple comparison correction (see fig. S3A-C for details). We nevertheless wondered whether pointing performance would be impacted by the location of the trajectories with respect to the clover axes, i.e. whether spatial updating for trajectories located on clover axes (dark gray trajectories on Fig. 3E, right) would be any better than for trajectories located off clover axes (light gray trajectories on Fig. 3E, right). We found a small but significant effect in which participants were better at pointing back at the object from an allocentric perspective when trajectories were situated on rather than off the clover axes (allocentric condition: 𝑡_(46)_ = −2.69, *p* = 0.009; egocentric condition: 𝑡_(46)_ = −1.1, *p* = 0.27, data averaged over bins 1/3/5/7 and 2/4/6/8, Fig. 3J).

## Rodent retrosplenial neurons exhibit clover-like tuning

We next asked whether the clover-like pattern of activity observed in humans also existed in the rodent brain, either at the individual cell or population level, and whether this spatial code was related to cells having vectorial attributes. To do so, we recorded single units from the rodent retrosplenial cortex (areas 29 and 30, deep layers, Fig. 4A and fig. S4A) during an open field foraging task in a rectangular arena (1.5m*1.5m) with a cue card on the north wall and a single circular intramaze object in the north-east quadrant (Fig. 4B). Analogous to human fMRI data analyses, we used GLM-based forward model selection to classify cells according to their vectorial allocentric and egocentric attributes (i.e., direction, distance, and their first derivatives) and combinations thereof (see details in Methods). Our analysis revealed that approximately 26% of the 149 single units exhibited allocentric directional and distance tuning (Fig. 4C, green bar), which was the most common functional classification among all covariates tested.

**Figure 4:**
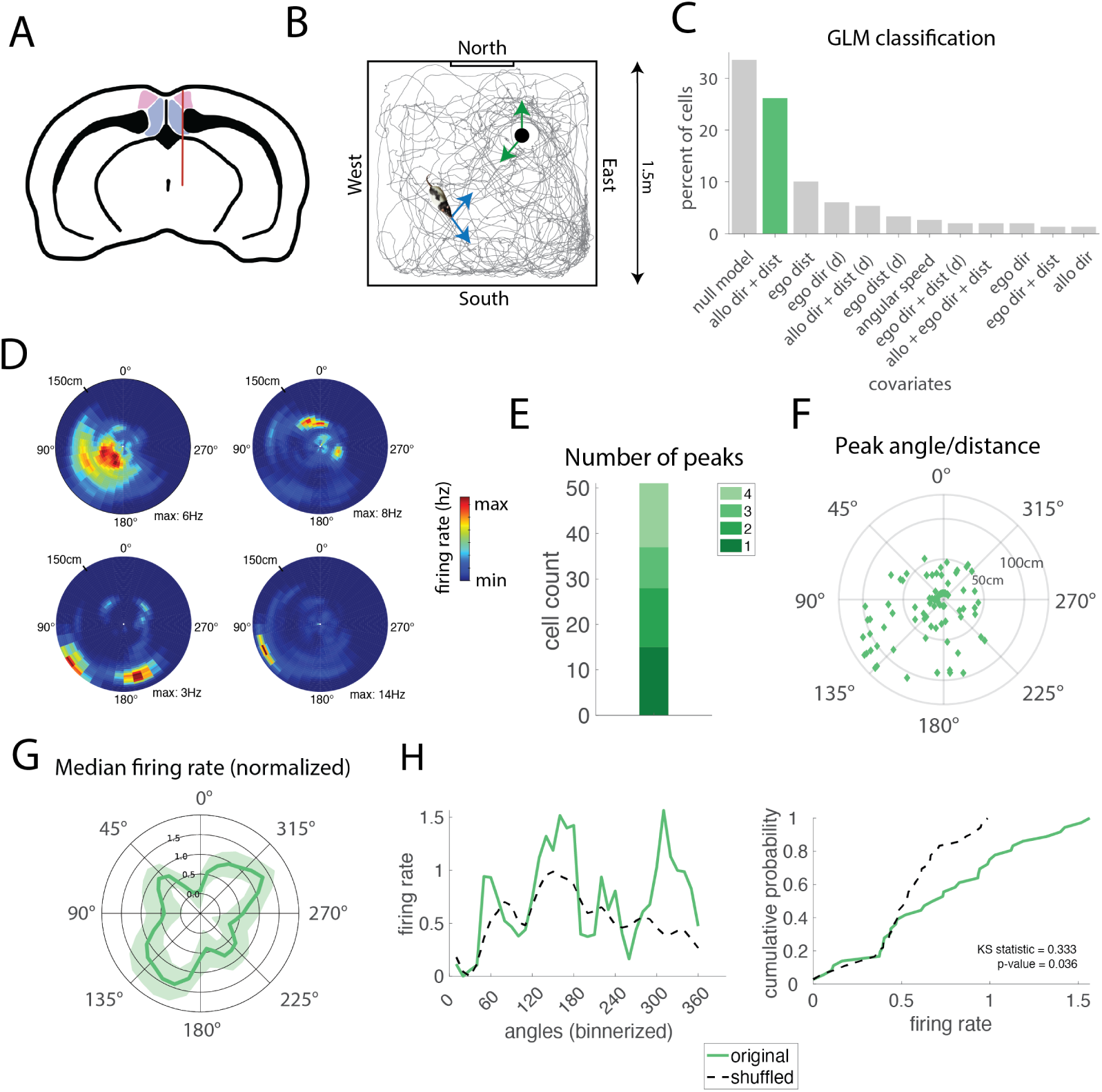
Allocentric vectorial coding in rodent retrosplenial cortex. (A) Recording sites in deep layers of areas 29 (blue) and 30 (pink) of the retrosplenial cortex; probe track shown in red. (B) Experimental arena (1.5×1.5 m) with cue card and object (black circle, NE quadrant); green and blue arrows indicate object-centric and egocentric frames. Animal trajectory shown. (C) GLM-based classification of neurons by explanatory covariates; green bar shows allocentric direction and distance tuning. Only covariates with ≥ 2 classified cells are shown. (D) Firing rate maps of four neurons with object-vector responses. (E) Distribution of firing rate peak counts per neuron. (F) Polar plot of peak locations (angle, distance) relative to object; Each dot represents a single peak. (G) Median normalized firing rate across neurons with allocentric tuning; Shaded area represents SEM. (H) Firing rate comparison: original (green) vs. shuffled (dashed); left: angle tuning; right: cumulative differences. KS = one-sided Kolmogorov-Smirnov test.

The spatial firing patterns of these neurons showed a notable structure, with individual cells displaying distinct firing fields around the object (Fig. 4D), the majority of which exhibited multiple peaks (Fig. 4E). The distribution of peak locations was evenly distributed across the population (Fig. 4F), with a notable clustering of peaks closer to the object. The population-level analysis revealed a clear clover-like pattern in the median normalized firing rates of cells (Fig. 4H), reminiscent of the activity patterns observed in humans. Statistical comparison between the original and shuffled data (Fig. 4I) confirmed the significance of this spatial organization (𝐾𝑆𝑠𝑡𝑎𝑡𝑖𝑠𝑡𝑖𝑐 = 0.333, *p* = 0.036), with the highest peak located approximately 45° off the north of the arena. By comparison, the same analysis performed on neurons with egocentric tuning properties (distance and direction relative to the animal) did not reach significance (𝐾𝑆𝑠𝑡𝑎𝑡𝑖𝑠𝑡𝑖𝑐 = 0.306, *p* = 0.069, see fig. S4B), suggesting that the clover-like pattern was a feature of allocentric vectorial cells in the recordings.

## Mnemonic rather than perceptual distance tuning

To estimate distance modulation on the human brain data, we normalized distance data with minimum/maximum distance as 1/-1. We found four peaks in our areas of interest: in RSC (coordinates: [-18,-59,12], negative modulation with increasing activity for increasing distance, Brodmann area 23), PHC (coordinates: [18,-35,-14], negative modulation, Brodmann area 36), EC (coordinates: [-24,5,-38], positive modulation) and HP (coordinates: [-22,-11,-12], positive modulation, Fig. 5A). The two medial temporal peaks were observed at uncorrected whole brain threshold only. We then discretized this data into 8 equidistant bins (Fig. 5B), but due to the nature of the task (start and end points of the trajectories are scattered around the object), middle bins corresponding to medium distance were observed more often than extreme bins, corresponding to very small or very large distances, leading to an unbalanced model (Fig. 5C). When estimating this model, we found a significant omnibus effect for the distance bins in RSC (𝐹_(2.95,94.40)_ = 5.56, *p* = 0.002, 𝜂_g_^2^ = 0.09) and PHC (𝐹_(2.84,85.28)_ = 2.95, *p* = 0.04, 𝜂_g_^2^ = 0.065), but no effect in EC (𝐹_(3.13,81.32)_ = 1.62, *p* = 0.19, 𝜂_g_^2^ = 0.036) or HP (𝐹_(3.82,76.48)_ = 1.10, *p* = 0.36, 𝜂_g_^2^ = 0.03). Proceeding with polynomial post-hoc contrasts we found a significant linear trend both in RSC (linear: 𝑡_(32)_ = 5.81, *p* < 0.0001; quadratic: 𝑡_(32)_ = −0.75, *p* = 0.45; cubic: 𝑡_(32)_ = 1.73, *p* = 0.09; quartic:𝐹_(32)_ = −0.96, *p* = 0.34; 14 outliers removed) and PHC (linear: 𝑡_(30)_ = 4.39, *p* = 0.0001; quadratic: 𝑡_(30)_ = −1.45, *p* = 0.15; cubic: 𝑡_(30)_ = 1.31, *p* = 0.19; quartic: 𝑡_(30)_ = −0.20, *p* = 0.83; 16 outliers removed), indicating a linear relation between distance and neural activity in those two clusters (Fig. 5D, see also individual examples in Fig. 5F).

**Figure 5:**
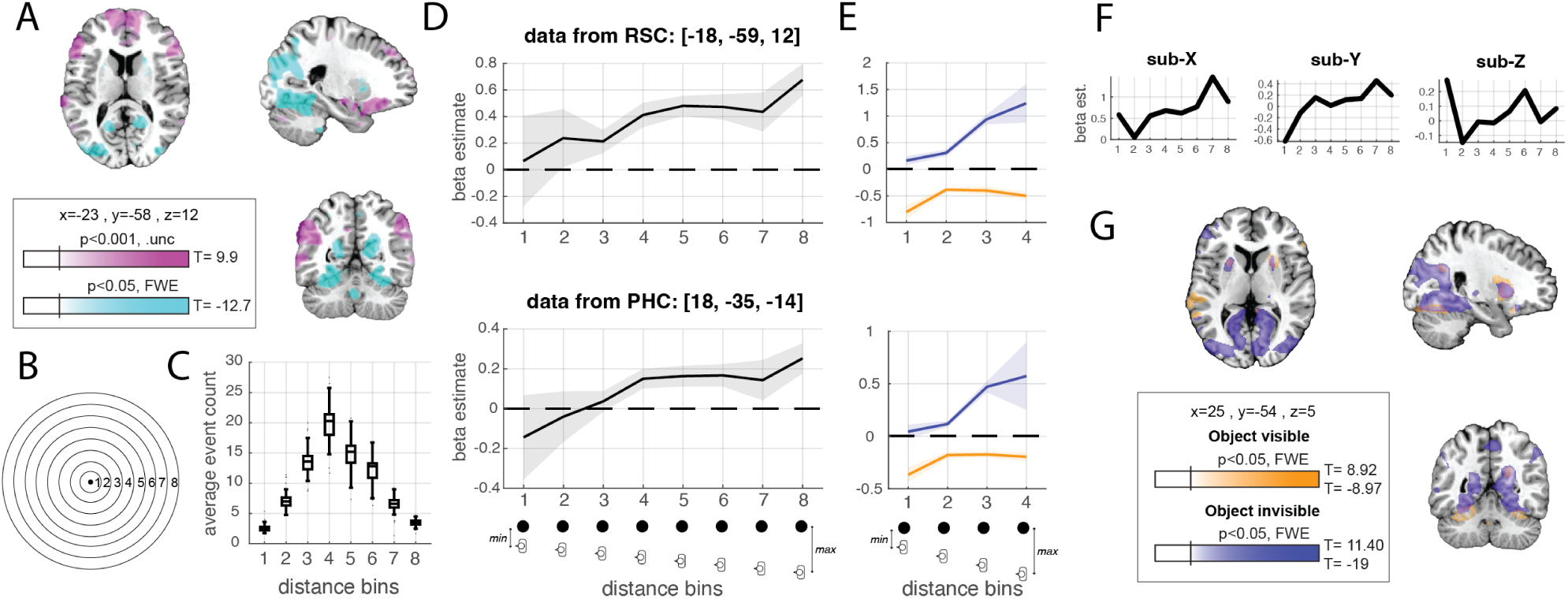
Distance modulation of the fMRI signal. (A) Localization model: whole brain parametric modulation with positive/negative modulation in pink/blue, respectively. Four peaks identified: retrosplenial cortex (RSC), parahippocampal cortex (PHC), entorhinal cortex (EC, anterior portion) and hippocampus (HP). Maximum, minimum T-scores and MNI slice location are presented. FWE=family-wise error correction. The EC and HP peaks were observed at uncorrected threshold only (unc.). (B-C) Distances used to estimate the tuning model. (C) Number of events for each bin with median, 25th and 75th percentiles (black box), 1.5 x interquartile range (whiskers), and outliers (dots). Note that this model is unbalanced. (D) Median tuning curves. Shaded areas show SEM. EC and HP tuning models were not estimated since the omnibus effect in these clusters did not reach significance (see text). (E) Median tuning curves in more parsimonious models with 4 bins, when the object was directly visible (yellow curves) or not (blue curves). (F) Individual tuning curves shown as best examples. (G) Whole brain parametric modulation with distance when object is visible (yellow) and invisible (blue). Negative and positive modulations are aggregated.

Based on the observation that the between- (Fig. 5D) and within-subject (Fig. 5F) variance in beta estimates was particularly high when participants came closer to the object (bins 1 and 2), we reasoned that those two areas could be sensitive to fast-changing visual inputs. We thus decided to rerun the model separating moments in time where the object was visible or invisible, and assess whether the distance modulation held. We found that RSC and PHC activity scaled up with increasing distance only when the object was invisible (Fig. 5G). We confirmed this observation with our tuning model (Fig. 5E), which accounted for the fact that these models were estimated on roughly half the amount data since the the number of distance bins was reduced to 4 (see Stastistics). Both areas indeed seemed to react to the immediate vicinity of the object, as the beta estimate in bin 1 significantly differed from the rest of the bins in the visible model (posthoc contrast: RSC: 𝑡_(29)_ = −4.14, *p* = 0.0003, 17 outliers removed; PHC: 𝑡_(39)_ = −2.88, *p* = 0.006, 7 outliers removed). However, past bin 1, RSC and PHC still coded for distance when the object was out of sight (invisible model in RSC: linear: 𝑡_(40)_ = 8.72, *p* < .0001, quadratic: 𝑡_(40)_ = 2.71, *p* = 0.02, does survive Bonferroni correction, 6 outliers removed; invisible model in PHC: linear: 𝑡_(39)_ = 9.52, *p* < .0001, quadratic: 𝑡_(39)_ = 3.48, *p* = 0.0025, 7 outliers removed) but not when the object was visible (visible model in RSC: linear: 𝑡_(29)_ = 0.67, *p* = 1, quadratic: 𝑡_(29)_ = 0.79, *p* = 0.87, 17 outliers removed, visible model in PHC: linear: 𝑡_(39)_ = 1.01, *p* = 0.63, quadratic: 𝑡_(39)_ = 0.58, *p* = 1, 7 outliers removed). No significant association was found between distance modulation and task performance.

## Topographical gradients of egocentric/allocentric modulation

Beyond examining peak location and tuning curves within our fMRI regions of interest, we next aimed to describe more specifically the anatomical topography of those responses. To do so, we averaged the parametric modulation maps across participants (fig. S5A-I). The pattern in RSC for the egocentric direction model, although peaking in Brodmann area (BA) 23, extended medially and ventrally within the RSC complex to BA 29 and 30 (fig. S5A-C), whereas the allocentric direction and distance models both yielded higher activation in the dorsal-lateral RSC, near BA 23 and 31. Regarding PHC, stronger modulation was observed in the posterior-medial portion of the ROI, with no clear difference between the three models. In EC, the allocentric direction modulation mapped posterior-medially, corresponding to the pmEC subregion. Finally, HP and EC showed signs of distance modulation in the anterior sections of both regions.

## Discussion

Using a novel spatial updating task and high-resolution 7T MRI in humans, we describe key navigation-related brain areas (e.g. retrosplenial, parahippocampal and entorhinal cortex) that dynamically represent vectorial information (distance and direction) to a proximal object, with specific activity patterns for egocentric/self-centered and allocentric/object-centered responses. On the one hand, egocentric signals peaked when the object was behind the navigator, while distance signals emerged only when the object was out of view. This suggests these signals may act as mnemonic buffers, helping the brain maintain representations of important objects even when they are no longer visible — enabling vision-independent spatial mapping. On the other hand, allocentric signals formed a four-direction clover-like pattern that aligned in most human participants on visually distinctive features in the environment. Parallel electrophysiological recordings revealed a similar four peak pattern of activity in the rat retrosplenial cortex, with allocentric vectorial cells contributing to the emergence of the clover pattern. In humans, the concurrent spatial updating task provided behavioral readouts of participants’ ability to track the object while moving around the environment. We found that one’s ability to keep track of their allocentric positioning relative objects was more accurate along the clover axes, highlighting this spatial code functional relevance for navigation in humans.

Brain activity clustered around four allocentric directions in retrosplenial (RSC), parahippocampal (PHC), and entorhinal cortex (EC). What is the possible origin and nature of this pattern of activity? First, we do not believe that the clover pattern is simply a reflection of the environmental layout, as was the case for travel axis-tuned neurons in subiculum (*16*) or multidirectional head direction cells in retrosplenial cortex (*17*), where bidirectional or quadridirectional firing was controlled by boundary arrangements or compartment symmetries, and disappeared in an open arena. Similarly in humans, the six-fold symmetry generally observed in the entorhinal cortex (*18*) can be transformed into a four-fold symmetry by local barriers (*19*). The fact that the clover was observed in environments with simple square walls (rodent experiment) or no walls at all (human experiment) points towards an independence from boundary layout. Similarly, this clover-like pattern seemed independent of landmark *structure*, as the arena used for humans contained multiple distal landmarks whose arrangements did not match the clover axes organization. However, the consistent alignment of the clover axes among different human participants clearly indicated that the clover anchoring was dictated by distal landmarks, likely the most salient one accessible. The clover axes in rodents were also tilted away from the Northern cue card, but the underlying principle organizing the anchoring of the clover axes (33.75^◦^ from the main landmark in humans or 45^◦^ from the cue card in rodent) needs further investigation. Second, the clover-like pattern was task-independent, as it was observed in complex tasks in humans and a simple free foraging tasks in rodents. This suggests the clover would be present at all times to inform and support object-self representations. This direct link to spatial cognition is demonstrated by improved allocentric spatial updating when participants navigated along the clover axes. Further research is needed to understand how these object-based representations interact with boundary-based representations (*20,21*) to facilitate goal-directed behavior, as seen in humans (*14, 15*) or rodents (*22*). We suggest the localization maps described here in humans could be used as translational basis for informing future targeted studies in rodents. Finally, if it would theoretically be beneficial for all distances and directions to be mapped evenly, the emergence of four peaks in the averaged firing rate of allocentric vector cells suggests an additional population code, akin to a neural reference axis that anchors object representations. The orientation of this neural reference axis happens to provide better object representation at a behavioral level, as shown in the human task. Collectively, our results support the idea that the four peak structure is a reflection of vectorial codes in the brain and is expressed in a population code that the summative nature of the BOLD signal is able to detect.

Brain activity varied based on the egocentric direction to an object, following a quadratic shape, with the highest activity in retrosplenial cortex (RSC) and parahippocampal cortex (PHC) when the object was positioned behind the participant. This suggests that these areas may compensate for the lack of direct visual experience of the object in order to maintain its direction representation. The strength of this quadratic modulation in the posterior parietal cortex (PPC) was reversed (higher beta weights when the object is in front), while RSC activity showed contralateral bias, with the left and right hemispheres representing contralateral object directions, consistent with previous findings in rodents (*3*). Finally, distance modulation was observed in human RSC and PHC. Activity in those brain areas was strongly influenced by the immediate vicinity of the object, and our control analysis indicated that PHC and RSC appeared to code for distance only when the object was not in sight. It is possible that visual inputs solely (e.g. monocular and binocular depth perception, or motion parallax) are sufficient to estimate distance to the object when it is visible (*23*). This information processed in visual cortex might inhibit or saturate activity in RSC/PHC, which then would act as memory buffers to maintain the representation of the object only when it cannot be perceived directly. It is an exciting possibility that the mnemonic nature of this direction/distance codes could extend beyond spatial navigation to more abstract cognitive processes such as episodic memory or time perception.

In summary, by using parallel neuroimaging in humans and electrophysiology in rodents, our study provides the first direct evidence of vector-based spatial coding in the human brain, which is mirrored in rodent vector cell activity, suggesting a conserved and potentially universal principle of neural representation that supports spatial navigation and vision-independent space mapping. We believe these vectorial brain mechanisms complement spatial codes for location (*24*), path integration (*11*) and heading (*12*) by providing a neural reference axis that anchors object representations and permit flexible navigation, while being particularly advantageous in environments containing proximal objects but lacking natural boundaries. It remains to be determined if and how these spatial codes are used as a basis for vector-based, goal-oriented navigation (*22*) and possibly cognition beyond space more generally (*25, 26*).

## Acknowledgments

The authors wish to warmly thank Annika E. Sauter for her critical contribution during the piloting phase of this study. This work was funded by the K. G. Jebsen Foundation, The Liaison Committee for Education, Research and Innovation in Central Norway (Samarbeidsorganet), and St. Olavs Hospital Trondheim. CD’s research is supported by the Max Planck Society, the European Research Council (ERC-CoG GEOCOG, grant No. 724836), the Kavli Foundation, the Jebsen Foundation, Helse Midt Norge and The Research Council of Norway (grants No. 223262/F50; 197467/F50). JRW’s research is funded by the Research Council of Norway FRIPRO (grant No. 300709), the Centre of Excellence scheme of the Research Council of Norway (Centre for Algorithms in the Cortex, grant No. 332640), the National Infrastructure scheme of the Research Council of Norway – NORBRAIN (grant No. 197467) and the Kavli Foundation.

## Author contributions

M.B., I.M.K. and C.F.D. designed the human experiments. M.B., P.S. and J.R.W. designed the rodent experiments. M.B., I.M.K. and P.S. collected and analyzed the data. M.B., I.M.K., P.S., J.R.W., C.F.D. wrote the article.

## Data and materials availability

The fMRI data supporting the findings of this study were collected in accordance with Norwegian data protection and privacy regulations. Due to legal and ethical restrictions imposed by the Norwegian Data Protection Authority (Datatilsynet) and the Regional Committees for Medical and Health Research Ethics (REK), the raw imaging data cannot be shared publicly. Data access is therefore limited to approved research collaborators who have obtained the necessary ethical approvals and data handling agreements in Norway. Interested researchers may contact the corresponding author to discuss potential collaborations under these constraints.

Additionally, group-level brain maps and individual tuning curves supporting Figs. 2,3,5,S5, as well as behavioral data supporting Fig. 1 (position and eye movement time series, task performance) and the raw rodent data supporting Fig. 4 will be made available on the following repository (https://osf.io/5t49x/) at the time of publication. The Matlab code used to model and analysis the fMRI data, as well as behavioral analyses scripts will be published alongside the data.

## Supplementary materials

Materials and Methods

Figs. S1 to S5

## Materials and Methods

### Human experiments

#### Participants

57 participants are enrolled in this study. Four participants are excluded due to nausea and six participants are excluded due to experimental error or scanner issue, leading to a final sample size of 47 (range: 18-34 years, 𝜇=25.6, 𝜎=4.12, 20 females, 27 males). All experimental procedures are performed in accordance with the tenets of the Declaration of Helsinki, and are approved by the Norwegian Regional Committee for Medical and Health Research Ethics (REK Midt, 17.02.2021–220248). All participation is voluntary. Potential participants are recruited by word of mouth or posters at the University and Hospital campuses. All participants give informed consent approval. 14 participants out of 47 are invited a second time to perform a control experiment (see Methods). Participants receive a financial compensation of 120 NOK (approximately 12 euros) per hour.

Inclusion criteria for this study are: (i) healthy, competent volunteers between the ages of 18 to 35, (ii) normal or corrected-to-normal vision, (iii) normal, uncorrected hearing, (iv) English fluency. Exclusion criteria are: (i) ongoing neurological or psychiatric treatment, (ii) history of brain injury, surgery or epilepsy, (iii) work in the metal-processing industry, (iv) pregnancy, (v) presence of foreign objects in the body, such as pacemaker, defibrillator, insulin pumps or metallic splinters, (vi) claustrophobia. MRI-compatible lenses are inserted on the head coil for participants requiring visual correction.

#### Task

The virtual environment is developed with Unity® game engine (Unity® Technologies, version: 2019.3.13f1, Fig. 1A-B). The environment is a grass-covered circular arena (radius: 300 virtual units [vu]). The arena is surrounded by water forming a natural, wall-free boundary. In the background, a mountain range with identifiable landmarks (eg. castle, lighthouse, church, windmill, trees, houses, rocks) is visible. The intramaze object, a column (dimension: 6.5*6.65*23.22 vu) is located 28.3 vu North-East off the arena center. The participant experiences a first-person view (camera height: 13.4 vu), with a 60-degree field of view.

The task consists of ten blocks of nine trials, alternating between egocentric and allocentric conditions, for a total of 90 trials (Fig. 1C-D). A single trial is composed of a self-paced free *navigation phase* to a starting location represented by a 4 vu-wide red circle on the arena ground, followed by the *orientation phase*, where participants get unlimited time to rotate in place to orient themselves around the starting location. After pressing the response button, the *trajectory phase* starts, where the participant is passively moved along a predefined path. Trajectories lie at 3 possible distances (”close”: 80 vu, ”medium”: 115 vu and ”far”: 160 vu) from the object and 3 possible directions to the object (”towards”: 60^◦^, ”perpendicular”: 90^◦^ and ”away”: 120^◦^ away from straight ahead direction. The order of the trajectories across different distances and directions are counterbalanced between blocks but not between participants. Each trajectory is experienced twice, once in the egocentric condition and once in the allocentric condition, to ensure a strictly similar visual experience between the two conditions. At the end of the trajectory and after a 2-second black screen (*black screen phase*), the participant has to point in the direction of the object, from an egocentric or allocentric perspective (*response phase*). In the egocentric condition, participants are asked to indicate where the object is relative to her/his own perspective after they had been moved along the trajectory (i.e., ahead, left, right or behind them), while in the allocentric condition, participants must indicate where they are relative to the object at the end of the trajectory (i.e.,North, East, West, or South of the object). During the response phase, two response panels can be displayed: an egocentric response panel, where the circle-like coordinate system is centered on the person, and an allocentric response panel centered on the column. Participants use the response buttons to rotate an arrow to indicate their response. The *feedback phase* follows where 5 different smiley faces are displayed according to pointing error (from happy to sad faces in the degree range: [0-15], [15-35], [35-65], [65-110] and [100-180]). The next trial starts immediately after. At the beginning of a new block, a panel indicated whether an allocentric or egocentric trial is to be performed. Participants are instructed during the out-of-scanner training that the main distal landmark (the castle) corresponds to the arbitrary North of the island. During the training, participants are shown a top-down view of the arena and its distal landmarks.

In the control experiment, the column height is reduced to 1.82 vu (its length and width did not change). Besides the column height, the environment and task used in the normal and control experiments are strictly identical.

#### Human protocol

The whole experiment lasts 2.5 hours on average. After consenting, participants perform a training task for up to 30 minutes outside of the scanner. After going through the 7T-MRI safety checklist with the on-site radiographer, the participant is set in the Siemens 7.0 T MAGNETOM Terra scanner (NTNU Norwegian 7T MR Center, PTX head coil). Participants are equipped with earplugs and foam pads for ear protection and minimizing motion. Throughout the whole session, participants are instructed to maintain complete stillness. Structural scans are acquired before functional ones. Functional scans are divided into 10-minute runs, with in-between breaks. Since the task is self-paced and in-scanner time is no longer than 1.5 hours, some participants perform fewer than the 90 predefined trials. An MR-compatible eye-tracker (SR Research EyeLink 1000) records eye movements when participants perform the task. A 5-point calibration procedure is performed after structural scans and prior to beginning the task. During the task, gaze position are queried at 60 Hz and integrated into the Unity environment using the EyeLink Wrapper 2.0 plugin package (*27*) when saccades are detected. Eye movements could not be recorded when participants wear corrective lenses, leading to a valid subset of n=37/47 participants with valid eye-tracking data. The virtual environment was projected on a 32 inches diagonal LCD screen sitting at the back on the bore (Nordic Neurolab InRoom device), seen through a mirror spanning approximately 30^◦^ of visual angle. The participants used four-button response grip controller (Nordic Neurolab ResponseGrips) to move in the virtual environment. Three buttons are used for movement: forward movement, left rotation, and right rotation, and one button to indicate their response. After a compulsory break, participants perform two cognitive tests before debriefing with the experimenters: a computerized version of the Perspective Taking test (*28*) and the 4 Mountains test (*29*) (4MT).

For participants invited twice, the order of the normal and control experiments is counterbalanced. Data from this second session is not presented here.

#### fMRI acquisition parameters

High-resolution T1-weighted images are acquired with an MP2RAGE sequence (Magnetization Prepared with 2 Rapid Gradient-Echoes), with following parameters: echo time=1.99 ms, repetition time=4.3 s, inversion time 1=840 ms, inversion time 2=2370 ms, voxel size=0.75 mm^3^, flip angle=5/6^◦^, slice thickness=0.75 mm, and a total of 224 sagittal slices.

BOLD T2*-weighted functional images are acquired with multi-band accelerated echo-planar sequence (*30*) and following parameters: TR=2 s, TE=19 ms, flip angle=80^◦^, voxel size=1.25 mm isotropic, field of view (FoV)=200 mm, 74 slices, GRAPPA multi-band acceleration factor=2, resulting in 288 volumes. Four volumes are acquired subsequently with reversed phase encoding direction for distortion correction.

### Data preprocessing

#### behavioral data

Trial outliers were identified and removed from the behavioral data. Outliers corresponded here to any pointing error value that is more than three scaled median absolute deviations from the median, subject-wise.

#### Gaze data

Participant’s 3D gaze data is smoothed using a 5 ms moving window. The intersection points between the participant’s gaze in Unity coordinates and the object, background, and ground are calculated using the MatGeom library. The colliders was a rectangular parallelepiped for the column (31*31*34.5 vu), a sphere for the background (radius: 300 vu), and a circular surface the ground (radius: 300 vu), centered on the arena terrain. Gaze positions from the orientation phase of the experiment only are used for calculating heatmaps. Heatmaps show intersections with the ground and the background only and are normalized across participants separately for the ego-centric and allocentric conditions. Additionally, gaze time proportion is calculated as the number of gaze intersections falling in each category (column, background, or ground) divided by the total number of recorded gaze intersections, separately for the egocentric and allocentric conditions.

#### Neuroimaging data

The T1-weighted background noise, typical of the MP2RAGE sequence, is removed by multiplying the uni and inv2 images (*31*). Subsequently, nifti-converted (*32*) (dcm2niix version 2-November-2020), BIDS-organized data were preprocessed with fmriprep (*33*) (version 23.0.2, RRID:SCR_016216), which is based on Nipype (*34,35*) 1.8.6 (RRID:SCR_002502).

#### B0 inhomogeneity mappings

The B0-nonuniformity map (or fieldmap) is estimated based on two (or more) echo-planar imaging (EPI) references with topup (*36*).

#### Anatomical data

The T1-weighted (T1w) image is corrected for intensity non-uniformity (INU) with N4BiasFieldCorrection (*37*), distributed with ANTs (*38*) 2.3.3 (RRID:SCR_004757), and used as T1w-reference throughout the workflow. The T1w-reference is then skull-stripped with a Nipype implementation of the antsBrainExtraction.sh workflow (from ANTs), using OASIS30ANTs as target template. Brain tissue segmentation of cerebrospinal fluid (CSF), white-matter (WM) and gray-matter (GM) is performed on the brain-extracted T1w using fast (*39*) (RRID:SCR_002823). Volume-based spatial normalization to one standard space is performed through nonlinear registration with antsRegistration (ANTs 2.3.3), using brain-extracted versions of both T1w reference and the T1w template. The following template is selected for spatial normalization and accessed with TemplateFlow (*40*) (23.0.0): ICBM 152 Nonlinear Asymmetrical template version 2009c (*41*) (RRID:SCR_008796; TemplateFlow ID: MNI152NLin2009cAsym).

#### Functional data

For each BOLD runs, the following preprocessing is performed. First, a reference volume and its skull-stripped version were generated by aligning and averaging 1 single-band references (SBRefs). Head-motion parameters with respect to the BOLD reference (transformation matrices, and six corresponding rotation and translation parameters) are estimated before any spatiotemporal filtering using Mcflirt (*42*). The estimated fieldmap is then aligned with rigid-registration to the target EPI (echo-planar imaging) reference run. The field coefficients are mapped on to the reference EPI using the transform. BOLD runs are slice-time corrected to 0.972 s (0.5 of slice acquisition range 0s-1.95s) using 3dTshift from AFNI (RRID:SCR_005927) (*43*). The BOLD reference is then co-registered to the T1w reference using mri-reg (FreeSurfer) followed by flirt (*42*) with the boundary-based registration (*44*) cost-function. Co-registration is configured with six degrees of freedom. First, a reference volume and its skull-stripped version are generated using a custom methodology of fMRIPrep. Several confounding time-series are calculated based on the preprocessed BOLD: framewise displacement (FD), DVARS and three region-wise global signals. FD was computed using two formulations following Power et al. (*45*) (absolute sum of relative motions) and Jenkinson et al. (*42*) (relative root mean square displacement between affines). FD and DVARS are calculated for each functional run, both using their implementations in Nipype (following the definitions by Power). The three global signals are extracted within the CSF, the WM, and the whole-brain masks. Additionally, a set of physiological regressors were extracted to allow for component-based noise correction (*46*) (CompCor). Principal components are estimated after high-pass filtering the preprocessed BOLD time-series (using a discrete cosine filter with 128s cut-off) for the two CompCor variants: temporal (tCompCor) and anatomical (aCompCor). tCompCor components are then calculated from the top 2% variable voxels within the brain mask. For aCompCor, three probabilistic masks (CSF, WM and combined CSF+WM) are generated in anatomical space. The implementation differs from that of Behzadi et al. in that instead of eroding the masks by 2 pixels on BOLD space, a mask of pixels that likely contain a volume fraction of GM is subtracted from the aCompCor masks. This mask is obtained by thresholding the corresponding partial volume map at 0.05, and it ensures components are not extracted from voxels containing a minimal fraction of GM. Finally, these masks are resampled into BOLD space and binarized by thresholding at 0.99 (as in the original implementation). Components are also calculated separately within the WM and CSF masks. For each CompCor decomposition, the k components with the largest singular values are retained, such that the retained components’ time series are sufficient to explain 50 percent of variance across the nuisance mask (CSF, WM, combined, or temporal). The remaining components are dropped from consideration. The head-motion estimates calculated in the correction step were also placed within the corresponding confounds file. The confound time series derived from head motion estimates and global signals were expanded with the inclusion of temporal derivatives and quadratic terms for each (*47*). Frames that exceeded a threshold of 0.5 mm FD or 1.5 standardized DVARS were annotated as motion outliers. Additional nuisance timeseries are calculated by means of principal components analysis of the signal found within a thin band (crown) of voxels around the edge of the brain, as proposed by (*48*). The BOLD time-series are resampled into standard space, generating a preprocessed BOLD run in MNI space. First, a reference volume and its skull-stripped version are generated using a custom methodology of fMRIPrep. All resamplings can be performed with a single interpolation step by composing all the pertinent transformations (i.e. head-motion transform matrices, susceptibility distortion correction when available, and co-registrations to anatomical and output spaces). Gridded (volumetric) resamplings are performed using antsApplyTransforms (ANTs), configured with Lanczos interpolation to minimize the smoothing effects of other kernels (*49*). Non-gridded (surface) resamplings are performed using mri-vol2surf (FreeSurfer). Many internal operations of fMRIprep use Nilearn (*50*) 0.9.1 (RRID:SCR_001362), mostly within the functional processing workflow. For more details of the pipeline, see the section corresponding to workflows in fMRIprep’s documentation.

#### Regions of interest definition

Four regions of interest (ROIs) are chosen *a priori* for analysis: i) the entorhinal cortex, including both posterior-medial (pmEC) and anterior-lateral (alEC) subregions, as defined in Syversen et al. (*51*), ii) the hippocampus, including both posterior and anterior subregions and iii) parahippocampal cortex. Both the hippocampus and parahippocampal cortex ROIS are generated from individual participants’ ASHS segmentation of the medial temporal lobe structures (*52*). Each mask is transformed from native space into MNI space and resampled to EPI resolution using FSL FLIRT. To create group masks, participant-specific masks in MNI space are averaged, thresholded at 50%, and binarized using fslmaths. Finally, iv) the retrosplenial complex, we used a mask from Ramanoel et al. (*53*), comprising both RSC proper (Brodmann areas 29/30) and functional RSC (Brodmann areas 23/31).

### fMRI analyses

#### Overview

All analyses are performed in SPM12. Images are high-pass filtered with a 180-s cutoff. Nuisance regressors for the first-level models include six realignment parameters (x,y,z, pitch, roll, yaw) and their derivatives and quadratic terms, in-scanner motion spikes (defined as any data point over a framewise displacement of 0.9), and the average signal within individual brain mask or anatomically-derived eroded CSF and WM masks. Data are spatially smoothed with a Gaussian full-width-at-half-maximum kernel of 8 mm. Out-of-brain voxels are excluded from the analyses.

#### Localization models

Stick functions sampled each second of the navigation phase similar to Spiers et al. (*54*), while parametric modulators for these regressors contained either *i)* the egocentric direction to the object in the range [0,180] degrees, *ii)* the allocentric direction to the object, with values in the range [0,360] degrees, and iii) distance-to-goal values, in the range [0,220] vu. Each direction/distance values were rescaled onto [1,-1] range, with 1 representing straight-ahead direction in the egocentric model, North direction in the allocentric model and closest distance to the object in the distance model.

#### Tuning models

Egocentric, allocentric and distance values during the navigation phase were discretized into 8 regressors, as shown in figures 2D-F, 3F and 5D. A fine-grained allocentric tuning model (figure 3G) was evaluated with allocentric directions being discretized into 16 regressors representing 16 allocentric direction bins. Note that outlier data points were removed before calculating group median in this fine-grained, thus more noisy, model.

#### Additional models

The hemispheric lateralization model (Figure 2H) was estimated with egocentric direction remapped onto [1,-1] range, representing right and left directions, respectively. The pure clover model (Figures 3A and S5G) was estimated based on 2 regressors representing moments in time where the participant location coincides with on-clover directions (bins 2, 4 ,6 and 8) or off-clover directions (bins 1, 3, 5 and 7 on Figure 3E). The visibility models (presented in Figure 5E,G) were estimated separately for moments in time where the object was visible to the subject, or not in sight. To account for the fact that these models are estimated on half as much data points compared to the original model, we choose to decrease the number of distance bins to 4.

#### Statistics for human data

Raw data were transformed with a Box–Cox transformation to achieve normality (*55*) and outlier data points (defined as lower/higher than 1.5 inter-quartile range of the 25^𝑡ℎ^ or 75^𝑡ℎ^ distribution percentile) were removed. Normality of the data is verified by visual inspection of Q–Q plots. For repeated measure ANOVAs, degrees of freedom of the factor (𝑑𝑓 _𝑓 𝑎𝑐𝑡𝑜𝑟_ = 𝑘−1) and the error (𝑑𝑓_𝑒𝑟𝑟𝑜𝑟_ = 𝑑𝑓 _𝑓 𝑎𝑐𝑡𝑜𝑟_ × 𝑑𝑓_𝑠𝑢𝑏 𝑗𝑒𝑐𝑡_) are presented, with (𝑑𝑓_𝑠𝑢𝑏 𝑗𝑒𝑐𝑡_ = 𝑁 − 1). Greenhouse-Geisser correction is applied when Mauchly’s Test for sphericity is significant, hence affecting the degrees of freedom of the omnibus tests reported in the text. The generalized eta-squared (ges, 𝜂_g_^2^) is used as as effect size. When appropriate, the four first polynomial contrasts were estimated, evaluating linear, quadratic, cubic and quartic tendencies in the data. When normality and homoscedasticity were dubious, Wilcoxon signed rank test was used to compare paired data. Effect size for non-parametric tests were calculated as a correlation coefficient 𝑟, as 𝑟 = 𝑧𝑠𝑐𝑜𝑟𝑒/√𝑛. Alpha level for statistical significance was set at *p* < 0.05. Posthoc tests were evaluated with a Bonferroni correction. Exact p-values are reported, until *p* < 0.0001. Statistical tests were one-sided.

### Rodent experiments

#### Animals and surgery

An adult female Long Evans rat, 3 months old, was used for single unit recordings. The animal was housed in an enriched cage with 3 other female litter mates. One week prior to surgery, the rat was moved to an individual cage (45 x 44 x 30 cm) to allow for acclimation and, following surgery, to prevent potential damage to the probe implant. The animal was housed in a temperature- and humidity-controlled facility and kept on a reversed 12-hour light–dark cycle. All experiments were conducted during the dark phase. The animal either maintained its baseline weight or experienced weight gain during the experiments and had ad libitum access to food and water, both before and during behavioral experiments. All experimental and surgical procedures were carried out in compliance with the Norwegian Welfare Act and the European Convention for the Protection of Vertebrate Animals used for Experimental and other Scientific Purposes.

During the surgical procedure, the animal was initially anesthetized in a ventilated Plexiglas chamber using 5% isoflurane vapor. Anesthesia levels were then adjusted to 1.0–2.5% isoflurane throughout the surgery to maintain an appropriate depth of anesthesia. Body temperature was kept at 37°C with a heating pad. After the animal was fully anesthetized, subcutaneous injections of analgesics were administered: Metacam (meloxicam; 2.5 mg/kg) and Temgesic (buprenorphine; 0.05 mg/kg). Additionally, local anesthesia with 0.5% Marcain was applied under the scalp before making the surgical incision. The surgical site on the skull was prepared by exposing the area and sterilizing it with 0.9% saline and 3% hydrogen peroxide. Using a high-speed dental drill with an 0.8 mm burr, openings were made in the skull for the placement of skull screws and craniotomies above retrosplenial cortex based on predetermined coordinates. To provide a stable electrical reference, a stainless steel bone-tapping screw was securely inserted into the skull, serving as the ground and electrical reference for subsequent recordings. A Neuropixels 1.0 probe (IMEC) was stereotactically inserted into the right hemisphere of the rat, with electrical contacts oriented toward lambda. The insertion targeted the tissue spanning from the retrosplenial cortex to the superior colliculus, based on coordinates relative to bregma: AP -5.6 mm and ML 0.5 mm. External reference and ground wires were attached to a skull screw positioned in the left hemisphere at AP -1 mm and ML +2 mm, and sealed with SCP03B Silver Conductive Paint Bottle from Elfa. The probe outside the brain was air-sealed using bead-sterilized Vaseline and enclosed within a custom-designed, 3D-printed black plastic housing. The implant was secured with Venusflow (Unident AS) cured under UV light and black-dyed dental cement to minimize light-induced electrical interference.

Postoperative care included administering subcutaneous fluids, analgesics, and allowing the animal to recover in a heated 37 ° C chamber for 1 to 2 hours prior to recording.

#### Recording and behavioral training

The animal was handled extensively for 3-5 weeks prior to the experiment to ensure acclimation to the experimenter. Following the handling period, the rat was habituated to a black 150*150*50 cm recording arena over several days before undergoing surgery and recording sessions. The recording environment included the arena itself, a white cue card positioned on the north wall (Fig. 4B), and a hardware setup located on the south wall. The recording environment was kept at consistent as possible throughout the experimental period.

The animal was introduced to the object and habituated previously so that it was familiar during the recording sessions. Neural data were collected during open-field foraging sessions (2 x 20 minutes each) in which the rat freely foraged for corn puff crumbs scattered on the arena floor by the experimenter. For the object session, the object was placed in the northeast quadrant of the box, and a similar protocol as the open field session was used (2 x 20 minute sessions). Each block of the task consisted of an open field session followed by the object session to ensure complete coverage of the arena in each condition.

#### Data analysis

In vivo electrophysiological hardware included Neuropixels 1.0 Probes (IMEC), a National Instruments PXIe-1071 chassis and a PXI-6133 I/O module for recording analog and digital input (*56*). The implanted probes were connected via a headstage circuit board and an interface cable positioned above the head of the animal. Excess cable was counterbalanced using elastic string to ensure that the animal could move freely throughout the experimental arena.

Neural data were recorded with SpikeGLX acquisition software (Janelia Research Campus). Amplifier gain settings were configured at 500× for recording spikes and 250× for local field potential (LFP). An external reference and an AP filter were set with a cutoff frequency of 300 Hz. Culster cutting and spike sorting was performed using Kilosort4 (*57*). During each recording session, signals were collected from Bank 1, mapping 385 channels.

#### GLM-based cell classification

To identify neurons with allocentric vectorial tuning to objects, we employed a Generalized Linear Model (GLM)-based statistical framework as described in detail in Mimica et al. (*58*). This framework was used to evaluate the neural responses of recorded cells to specific behavioral and spatial features. Neural spike data were modeled as a function of the position of the animal, allocentric object distance and angle to the animal (and their derivatives), egocentric distance and angle to the object from the animal (and their derivatives), and other kinematic features such as turning and running speed.

The model selection process involved three steps: *i) Feature selection:* features related to animal-object vector relationships were extracted from behavioral data using a combination of 3D motion capture analysis, as outlined in Mimica et al. (*58*). These features included allocentric position of the animal relative to the object (allocentric head angle, distance to the object and their 1st derivatives) and egocentric object-animal proximity (egocentric head angle and distance to the object calculated from the nose of the animal and their 1st derivatives) and angular speed. *ii) GLM fitting:* neural spike trains were modeled using a Bernoulli family GLM, where the probability of spiking was predicted based on the selected behavioral features. 10 fold cross-validation was applied to ensure robust model fitting and avoid overfitting; *iii) Model evaluation:* the goodness-of-fit for each GLM was assessed using metrics including deviance, pseudo-R2, and log-likelihood ratios. Neurons were classified as exhibiting allocentric vectorial tuning to the object if their firing rates showed significant modulation by allocentric object proximity features, as determined by the likelihood ratio test (*p* < 0.05).

#### Data processing

The set of neurons identified by the GLM as having allocentric vectorial tuning to the object had to meet 3 further criteria, such that cells with potentially spurious or non-specific responses were removed where possible. These criteria included: *i)* Goodness-of-Fit threshold: only neurons with a pseudo-R2 above a predefined threshold (0.1) were retained. *ii)* significant tuning: neurons had to exhibit statistically significant tuning curves for allocentric features to the object. *iii)* firing rates: to reduce the occurrence of spurious or nonspecific responses, neurons with mean firing rates below 2 Hz were excluded, as were neurons with mean rates above 20 Hz. Neurons with mean firing rates below 2 Hz often fire too sparsely for reliable tuning estimates, and those above 20 Hz can reflect overly high, potentially nonspecific activity. Restricting to 2–20 Hz helps reduce spurious tuning and ensures a more consistent physiological range.

#### Single neuron firing rate

Spiking rates were normalized by occupancy time in each spatial bin, with spatial and temporal binning of spiking activity as follows:

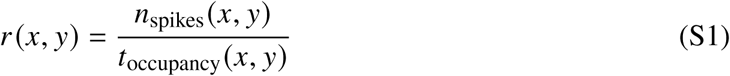

where 𝑟 (𝑥, 𝑦) is the firing rate in a specific bin defined by spatial coordinates (𝑥, 𝑦), 𝑛_spikes_(𝑥, 𝑦) is the number of spikes recorded in that bin, and 𝑡_occupancy_(𝑥, 𝑦) is the total time spent in that bin.

#### Population median firing rate across neurons

After calculating firing rate maps for individual neurons, we computed the median firing rate across all neurons at each bin location to summarize population activity:

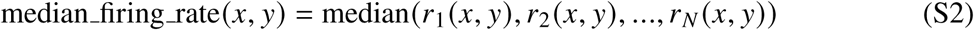

where 𝑟_𝑖_ (𝑥, 𝑦) represents the firing rate of neuron 𝑖 in bin (𝑥, 𝑦), and 𝑁 is the total number of neurons. This approach allows for a robust estimation of the central tendency of firing rates across the population and is less sensitive to outliers compared to the population mean. The standard error of the mean (SEM) was calculated to quantify variability in firing rates as follows:

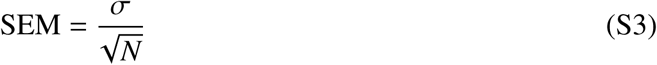

where 𝜎 is the standard deviation of the firing rates across cells, and 𝑁 is the total number of cells. For time points with missing data, the standard deviation was calculated excluding NaN values. Both the median firing rates and SEM were then smoothed with a Gaussian kernel:

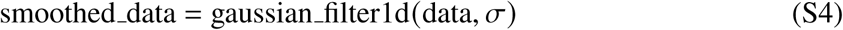

where 𝜎 = 0.75 for firing rates and 𝜎 = 0.1 for SEM.

#### Peak detection algorithm

The peak detection process involved the following steps: First, each neuron’s 2D firing rate matrix was normalized to ensure consistent detection across cells with varying firing rate ranges:

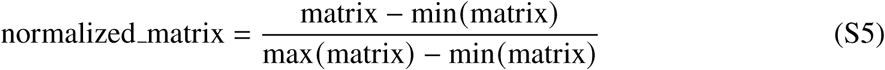

Second, a dynamic threshold was calculated for each neuron based on its maximum normalized firing rate:

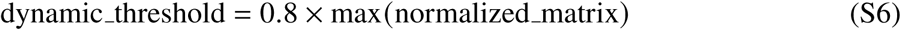

Third, the peak_local_max function from the skimage.feature module was used to detect local maxima in the normalized matrix. Parameters included the min_distance, set to 5 (in matrix indices) to ensure spatial separation between peaks, threshold_abs, set to the calculated dynamic threshold and the exclude_border, set to False to allow detection of peaks near matrix edges. Fourth, detected peaks were sorted by intensity in descending order, and the top 4 peaks were selected as significant. Finally, peak coordinates were converted from matrix indices to actual angle and distance values using predefined bin centers. This allowed for systematic identification of preferred firing directions and distances for each neuron, features relevant to their allocentric vectorial properties.

#### Statistics for rodent data

The Kolmogorov-Smirnov (K-S) test was used to compare the similarity between one distribution and a shuffled version of itself. We report the KS statistic, or D statistic, which is the maximum absolute difference between the cumulative distribution functions. The D value is a measure of the test effect size, as it represents how far apart the two distributions are (range from 0 to 1).

#### Histology

After the recordings were completed, the rat was euthanized with an overdose of isoflurane, followed by intracardial perfusion with saline and 4% paraformaldehyde. The probe shank was left in the brain and the entire skull was immersed in 4% paraformaldehyde for 24 hours. The next day, the brain was carefully extracted from the skull and placed in a 2% dimethyl sulfoxide (DMSO, VWR, USA) solution for cryoprotection, lasting one to two days before cryosectioning.

Following cryoprotection, the brain was frozen and sectioned coronally in three series of 40 𝜇m slices using a freezing sliding microtome (Microm HM-430, Thermo Scientific, Waltham, MA). The first series of sections was mounted directly onto Superfrost slides (Fisher Scientific, Goteborg, Sweden). Hoechst staining was performed to confirm anatomical probe placement (fig. S4A).

All brain sections were digitized using a digital scanner equipped with appropriate illumination wavelengths and scanning software provided by Carl Zeiss AS (Oslo, Norway). The resulting images were analyzed and visualized using ZEN (Blue Edition) software.

Probe placement and channel counts in cortical sub-regions were estimated using the HERBS (Histological E-data Registration in rodent Brain Spaces) software toolbox (*59*), which allowed for precise alignment of electrode tracks in histological sections and subsequent mapping of recording sites in each region.

---

**Figure S1:**
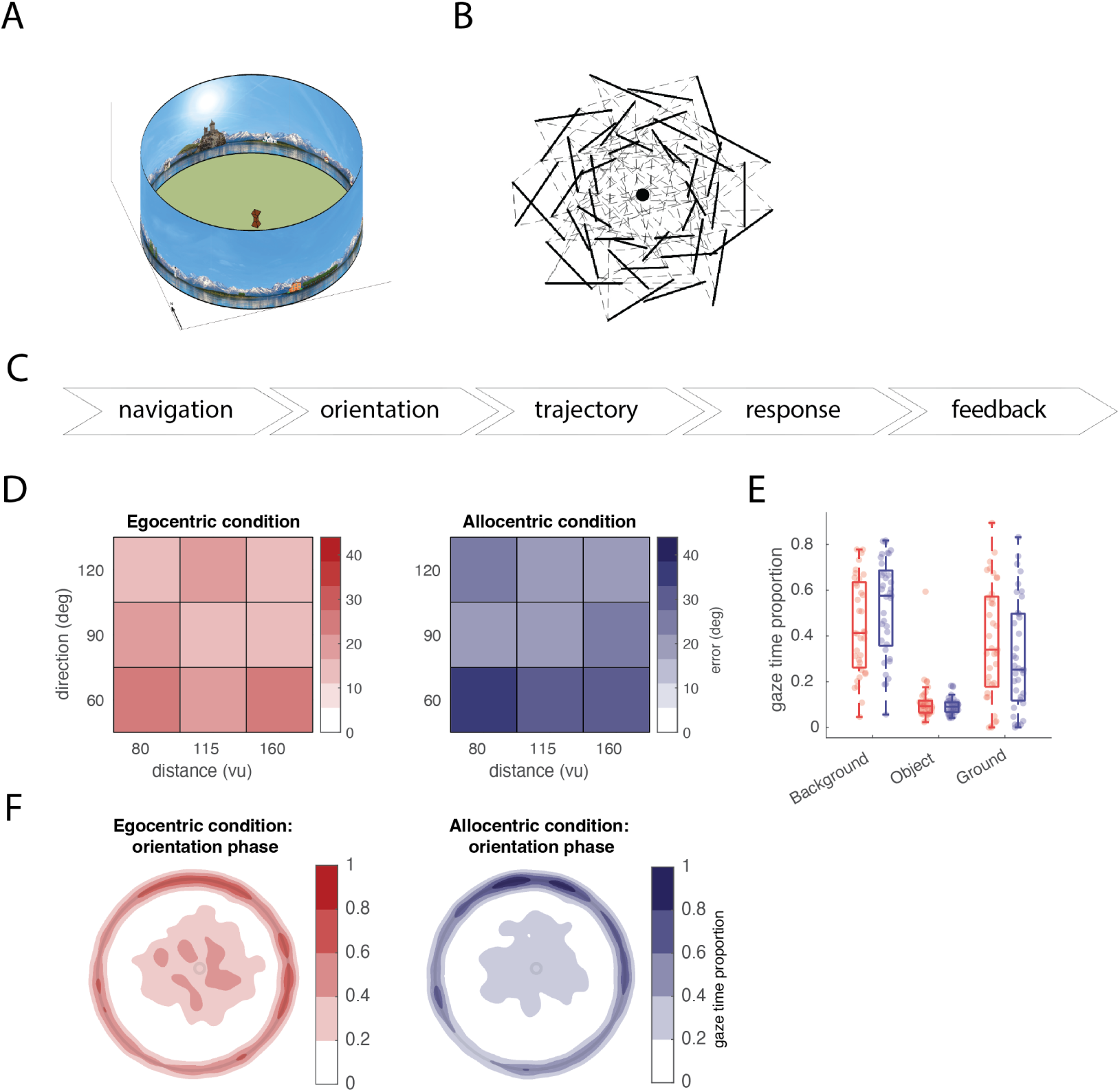
Extended behavioral spatial updating results. (A) Isometric view of the circular wall-free arena with an intramaze object (i.e. the column). (B) The task alternates between periods of active (dashed lines) and passive (thick lines) navigation, scattered around the object (black dot) to ensure full direction and distance coverage. (C) Task phases, as described in the main text. (D) Mean error in degrees across different distances and egocentric directions to the object. (E) Proportion of time spent looking at 3 areas of interests (the background, the object, the ground) during the orientation phase of the experiment, separately for the two conditions. (F) Gaze heatmaps during the orientation phase of egocentric and allocentric condition, respectively. Gaze intersections on the object were removed from the heatmaps for visualization purposes. Heatmaps were normalized collectively across the two conditions.

**Figure S2:**
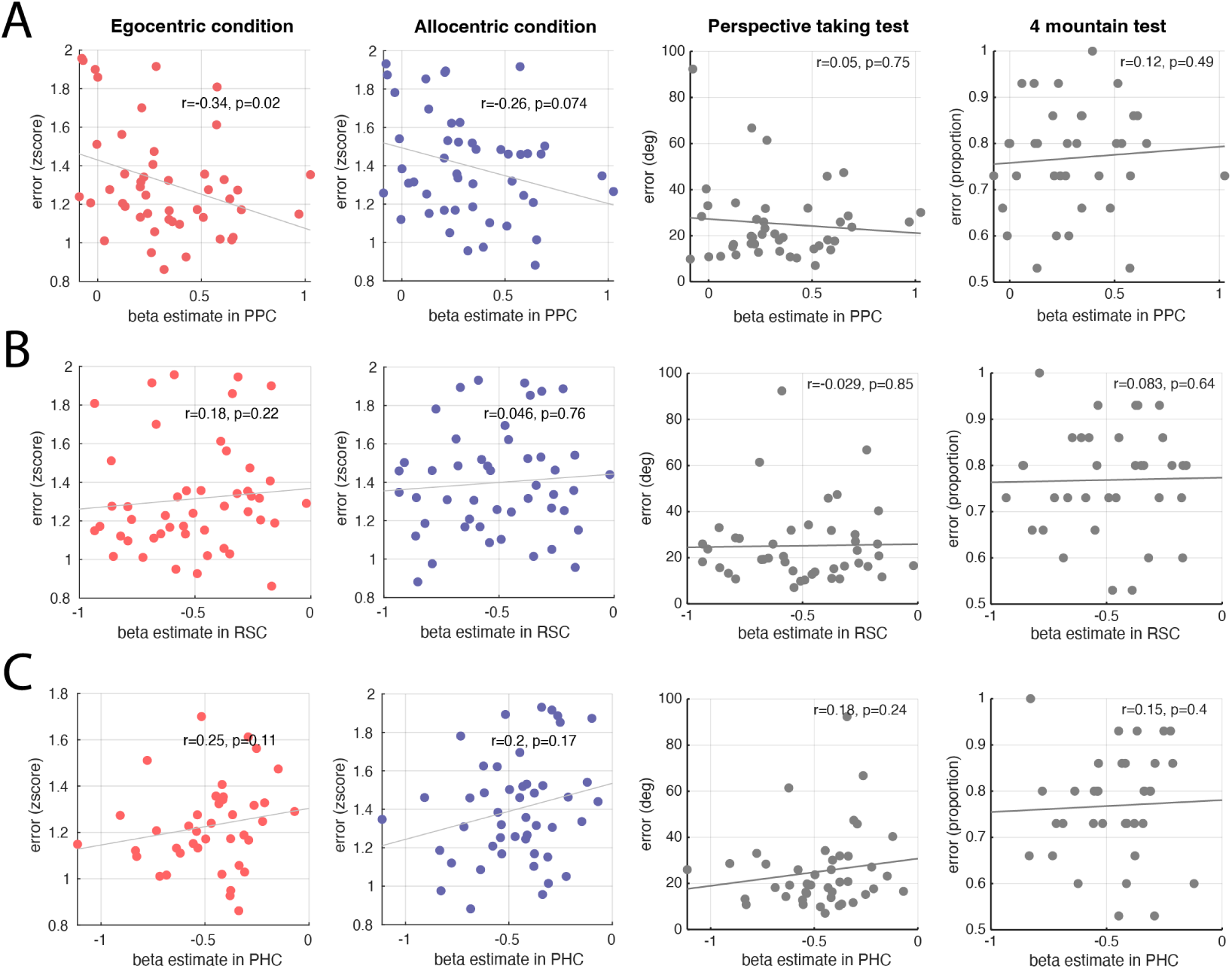
Correlation between the egocentric modulation strength, task performance and cognitive scores. Average beta estimate in PPC (A), RSC (B) and PHC (C) and z-scored error in the task, in the egocentric (left) and allocentric (middle-left) conditions and performance and the Perspective taking test (middle-right) and 4 mountain test (right). Rho and p-values are for Spearman correlations.

**Figure S3:**
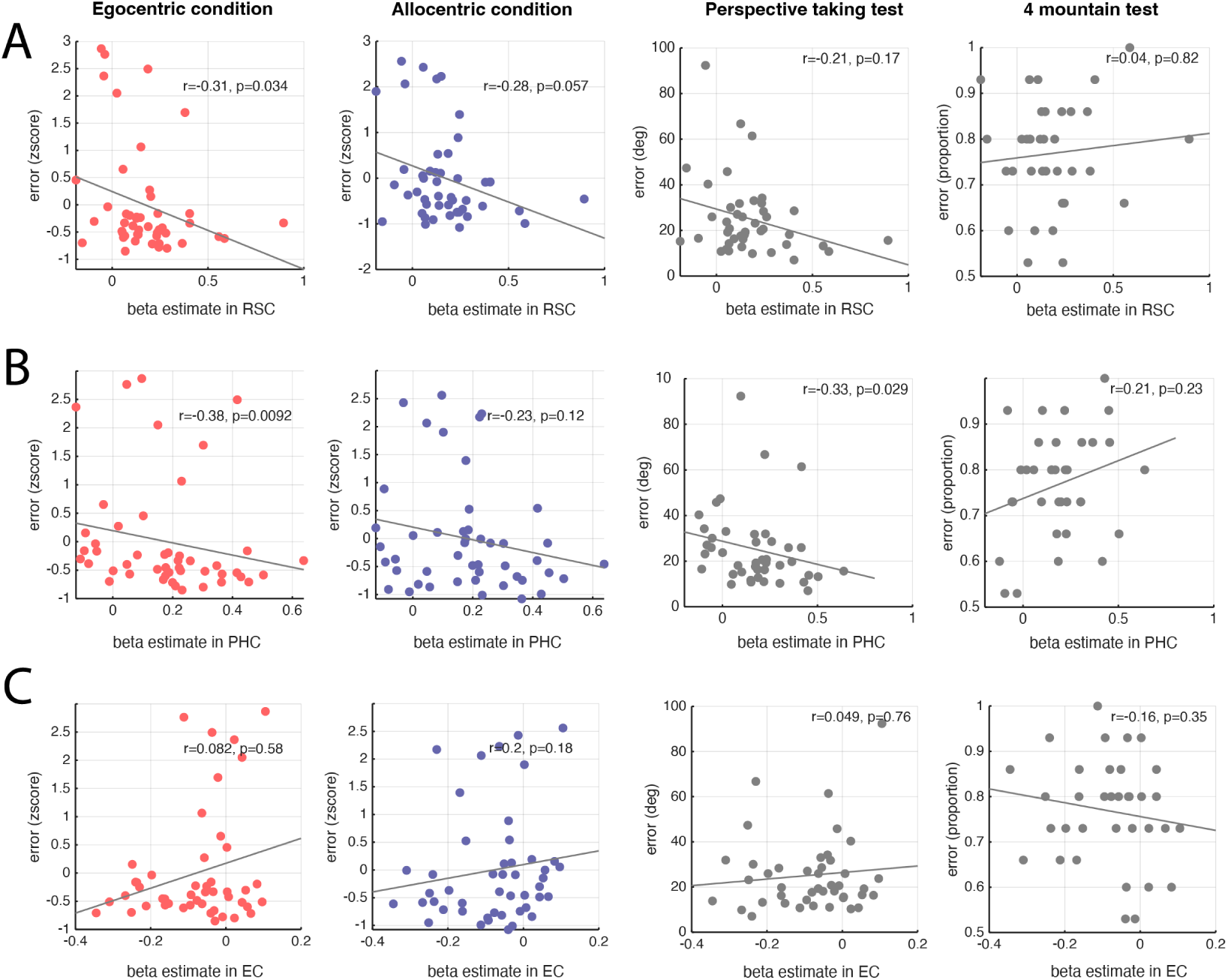
Correlation between the allocentric modulation strength, task performance and cognitive scores. Average beta estimate for the pure clover model in RSC (A), PHC (B) and EC (C) and z-scored error in the task, in the egocentric (left) and allocentric (middle-left) conditions and performance and the Perspective taking test (middle-right) and 4 mountain test (right). Rho and p-values are for Spearman correlations.

**Figure S4:**
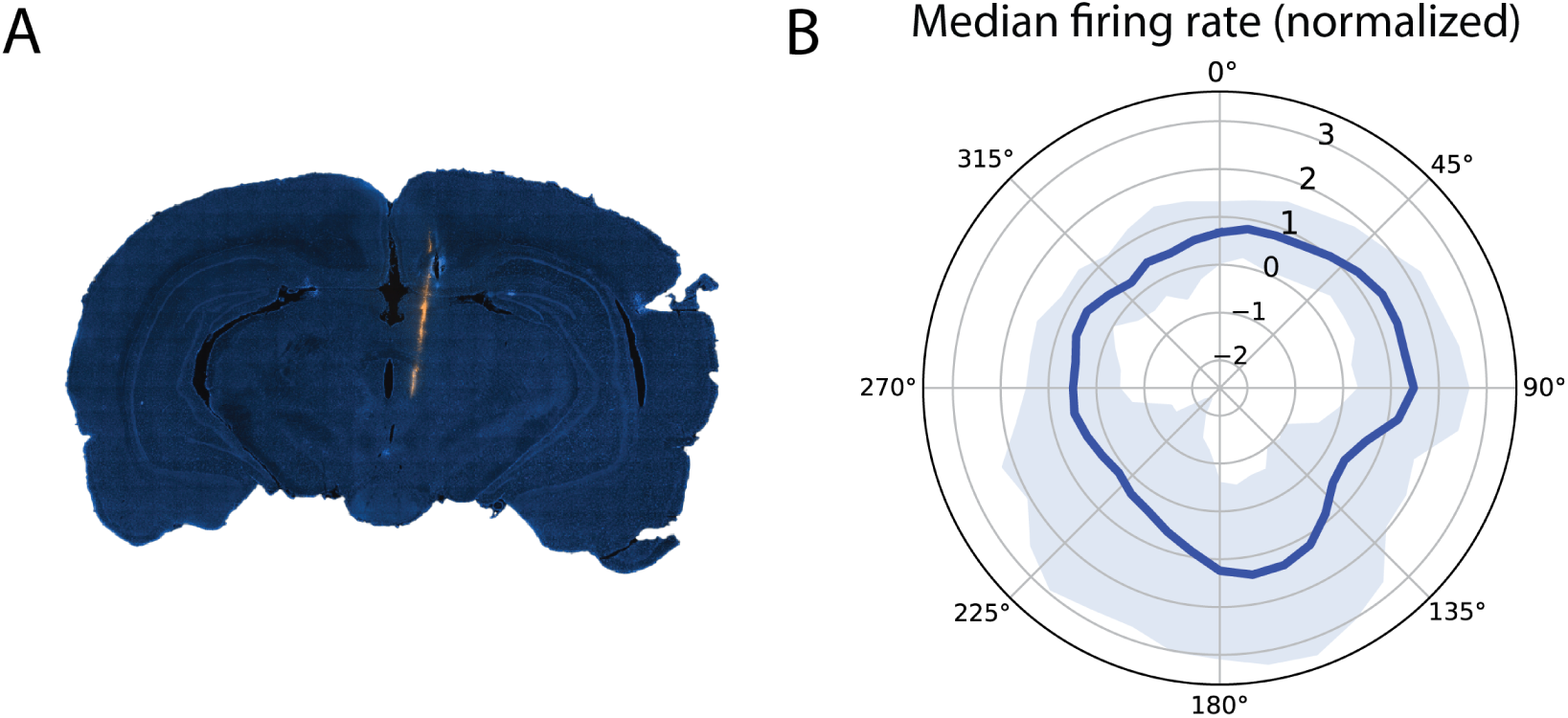
Extended electrophysiological results. (A) Histological verification of recording location in retrosplenial cortex area 29 and 20 deep layers, showing probe track in a coronal brain section stained with Hoechst. (B) Population analysis showing the median normalized firing rate across all neurons with egocentric vectorial properties. Blue line indicates the median and shaded area represents the standard error of the mean; no particular pattern emerged from the data.

**Figure S5:**
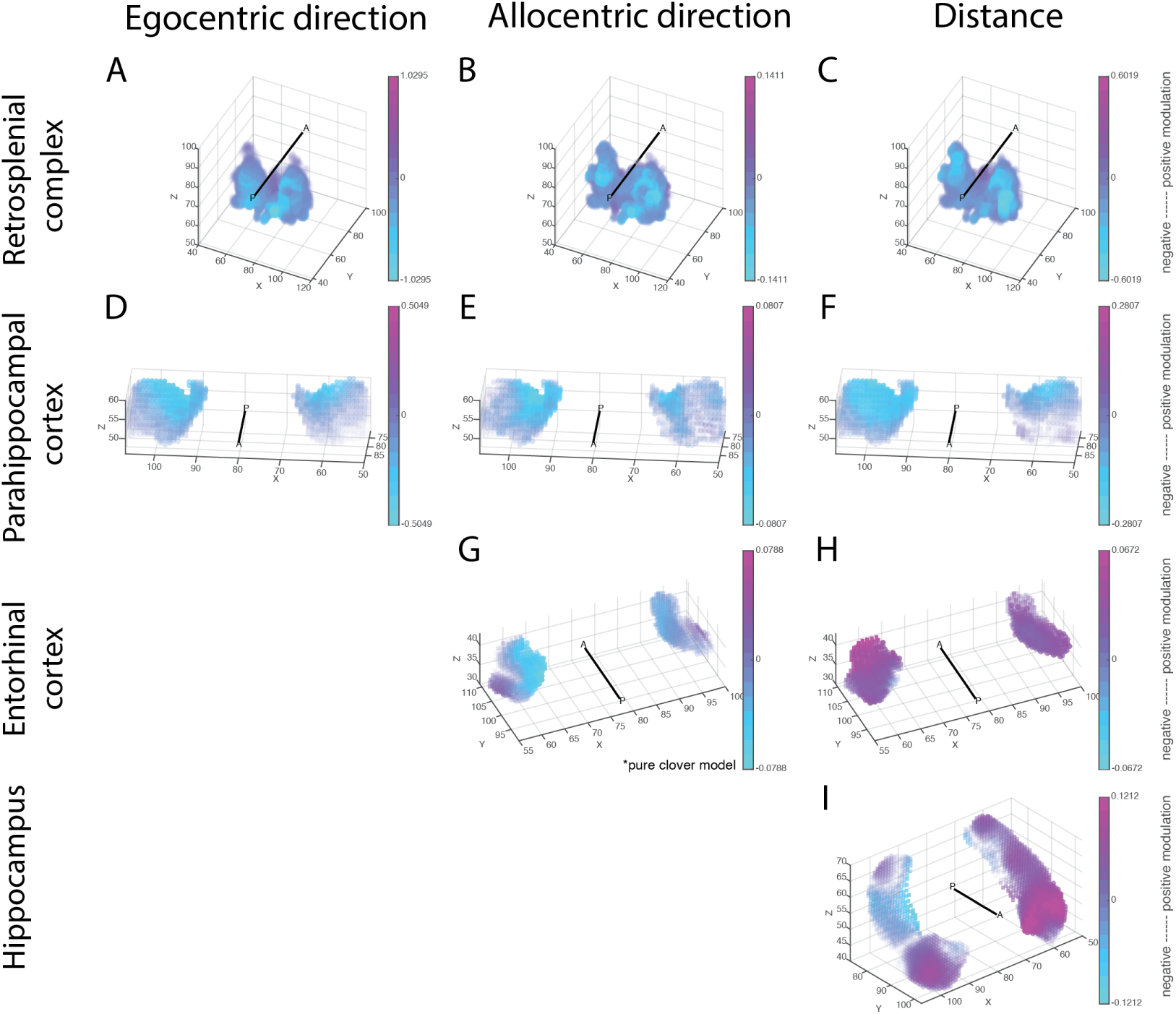
Topographical gradients of ego- and allocentric direction and distance modulation. (A-I) Each map displays the parametric modulation for each voxel in the ROIs, averaged across participants. 3D locations are expressed as voxel indices, with the anterior-posterior (A-P) axis shown on each panel. Color bars are normalized by the minimum and maximum values within a bilateral ROI. Additionally, a transparency map was added to the color map, with higher transparency values for parametric modulation values closer to zero.

